# Histone deacetylase inhibitors reduce the number of herpes simplex virus-1 genomes initiating expression in individual cells

**DOI:** 10.1101/086249

**Authors:** Shapira Lev, Ralph Maya, Tomer Enosh, Cohen Shai, Kobiler Oren

## Abstract

Although many viral particles can enter a single cell, the number of viral genomes per cell that establish infection is limited. However, mechanisms underlying this restriction were not explored in depth. For herpesviruses, one of the possible mechanisms suggested is chromatinization and silencing of the incoming genomes. To test this hypothesis, we followed infection with three herpes simplex virus 1 (HSV-1) fluorescence-expressing recombinants in the presence or absence of histone deacetylases inhibitors (HDACi’s). Unexpectedly, a lower number of viral genomes initiated expression in the presence of these inhibitors. This phenomenon was observed using several HDACi: Trichostatin A (TSA), Suberohydroxamic Acid (SBX), Valporic Acid (VPA) and Suberoylanilide Hydoxamic Acid (SAHA). We found that HDACi presence did not change the progeny outcome from the infected cells but did alter the kinetic of the infection. Different cell types (HFF, Vero and U2OS), which vary in their capability to activate intrinsic and innate immunity, show a cell specific basal average number of viral genomes establishing infection. Importantly, in all cell types, treatment with TSA reduced the number of viral genomes. ND10 nuclear bodies are known to interact with the incoming herpes genomes and repress viral replication. The viral immediate early protein, ICP0, is known to disassemble the ND10 bodies and to induce degradation of some of the host proteins in these domains. HDACi treated cells expressed higher levels of some of the host ND10 proteins (PML and ATRX), which may down regulate the number of viral genomes initiating expression per cell. Corroborating this hypothesis, infection with three HSV-1 recombinants carrying a deletion in the gene coding for ICP0, show a reduction in the number of genomes being expressed in U2OS cells. We suggest that alterations in the levels of host proteins involved in intrinsic antiviral defense may result in differences in the number of genomes that initiate expression.

## Introduction

Herpes simplex virus-1 (HSV-1) is a common human pathogen and is considered a prototype of the large herpesviridae family. Herpesviruses are a good example for viruses that coevolved with their hosts, to maintain a tight balance between minimal pathogenicity and maximal spread in the population. This balance can also be observed at the cellular level. Like many other double stranded DNA viruses, the viral genome replicates inside the host nucleus and utilizes host factors to facilitate viral replication. On the other hand, the intrinsic immunity of the cell evolved to inhibit the expression and replication of foreign DNA. Restriction factors are constitutively expressed host proteins that provide the first intracellular line of defense against viruses and other intracellular parasites (Bieniasz, 2004; Yan and Chen, 2012). Many mechanisms have evolved to recognize and inhibit viral DNA, including silencing the incoming genomes inside the nucleus of the infected cell.

Naked herpes viral DNA genomes enter the host nucleus through the nuclear pore complexes (Kobiler et al., 2012). Upon entry, the viral DNA is rapidly associated with host proteins inside the nucleus that can inhibit replication of the virus (Knipe, 2015). Among these proteins are the well characterized components of the nuclear domain 10 (ND10) bodies, also known as promyelocytic leukemia (PML) bodies (Maul et al., 1996). Some of these proteins including PML, hDaxx, ATRX, SP100 and more, were shown to have antiviral activity (Everett et al., 2006; Everett et al., 2008; Lukashchuk and Everett, 2010). ICP0, a viral immediate early protein with a E3 ligase activity interacts with these host proteins and facilitates their degradation, thus enhancing viral gene expression (Boutell and Everett, 2013). Other host proteins that interact with the naked viral DNA are histones. The viral DNA and histones form nucleosomes, which are known to be important for establishing either the lytic or the latent infection (Cliffe and Knipe, 2008; Oh and Fraser, 2008).

The host and viral chromatin state can be regulated by many post translational modifications of the histone proteins including: acetylation, methylation, phosphorylation, ubiquitination and more. The ‘histone code’ i.e. assigning functions to combinations of histone modifications is more complex than originally assumed (Jenuwein and Allis, 2001). However, the role of lysine acetylation of the histone tails is still mostly considered to be associated with open chromatin, a condition that favors transcription. Histone deacetylases (HDAC’s) are a group of enzymes that remove acetyl groups from an ε-N-acetyl lysine amino acid on a histone, allowing the histones to pack the DNA more tightly. An important role for the HDAC activity was identified in many key biological processes including development, cell proliferation, DNA repair and more. Inhibition of HDAC activity has a therapeutic value for many diseases including many malignancies. Therefore, many HDAC inhibitors (HDACi’s) were identified, with different specificities against the 18 known human HDAC’s (Falkenberg and Johnstone, 2014). HDACi were able to induce HSV-1 reactivation from latency (Arthur et al., 2001; Poon et al., 2003; Danaher et al., 2005; Neumann et al., 2007) and to enhance HSV-1 based oncolytic virus in some tumor derived cells (Otsuki et al., 2008; Cody et al., 2014). Recent findings suggest that different histone modifying factors are important for infection depending on cell type (Oh et al., 2014).

We previously described that using three isogeneic viral recombinants, each carrying a different fluorescent protein, we can estimate the number of viral genomes initiating expression (Kobiler et al., 2010; Kobiler et al., 2011; Taylor et al., 2012). The basis for our method is the assumption that as more genomes are being expressed per cell, more cells will express all three fluorescent colors. In brief, λ, representing the most likely average number of incoming viral genomes initiating expression per cell, was estimated given the distribution of cell colors in the cell population according to the following equation:

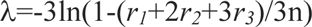

In which *r*_1_, *r*_2_, and *r*_3_ represent the numbers of cells expressing one-color, two-color, and three-color, respectively, and *n* represents the total number of colored cells that were analyzed (Kobiler et al., 2010). We found that during lytic infection, only a limited number of incoming herpes viral genomes can initiate expression and replication in a given cell (Kobiler et al., 2010; Kobiler et al., 2011; Taylor et al., 2012). Recently, we corroborated these findings with a single cell based method (Cohen E and O.K. unpublished), indicating that our mathematical model provides a good estimate for the number of viral genomes being replicated per cell.

We hypothesize that host factors alter the number of incoming genomes initiating expression and replication. We assumed that the histone modifying factors could be involved in this process. To test this hypothesis, we examined the role of HDACi during the initiation of gene expression by incoming herpes viral genomes. We found that treatment with HDACi results in a lower number of viral genomes that initiate replication per cell in different cell types. Treatment with HDACi results in increased levels of PML and ATRX, known intrinsic immunity proteins. Taken together, our results suggest that the level of host restriction factors modify the probability of a viral genome to initiate replication.

## Materials and Methods

### Cells

The experiments were performed with green monkey kidney cells (Vero cells, ATCC CCL-81), human immortalized foreskin fibroblasts (HFF cells) or human female osteosarcoma cells (U2OS cells ATCC HTB-96). The immortalized HFF cells were a kind gift from the Sara Selig. These HFF cells were immortalized by hTERT transfection. All cells were grown with Dulbecco's Modified Eagle Medium (DMEM X1; Gibco), supplemented with 10% Fetal Bovine Serum (FBS; Gibco) and 1% Penicillin (10,000 units/ml) and Streptomycin (10mg/ml) (Biological Industries Israel).

### Viruses

All viruses are derivatives of HSV-1 strain17+. Viral recombinants OK11, OK12 and OK22 carry a single fluorescent protein (mCherry, EYFP and mTurq2, respectively) with a nuclear localization tag under the CMV promoter between UL37 and UL38 genes as described previously (Taylor et al., 2012; Criddle et al., 2016). The fluorescence expressing, ICP0 deletion recombinants were constructed for this work. Shortly OK11 and OK22 were crossed with a viral recombinant with YPet protein inserted into the UL25 gene. Similarly, OK12 was crossed with a viral recombinant with mCherry protein inserted into the UL25 gene. These dual color viruses were further crossed with HSV-1 dl1403 strain. HSV-1 dl1403 strain (Stow and Stow, 1986), a 2kbp deletion in the each of the two copies of the ICP0 gene, was a kind gift from Roger Everett. The new cross selected for single color recombinants with ΔICP0 phenotype. Each cross was plaque purified to homogeneity on either Vero cells (for wild type recombinants) or U2OS strains (for ΔICP0 recombinants). Viral recombinants OK29, OK32 and OK40 carry a single fluorescent protein (mCherry, mTurq2 and EYFP, respectively) in an ICP0 deletion background. The new recombinants were tested by PCR, phenotype and growth curves (figure 5A and B). Viruses were grown and tittered on either Vero (for wild type recombinants) or U2OS (for ΔICP0 recombinants) cells. The multiplicity of infection (MOI) was calculated as the number of plaque forming units (PFU) per cell.

### HDACi protocol

Cells were seeded in 24 wells plates for 24hours at 37^o^C. Three hours prior to infection, the medium was replaced with medium containing the specific inhibitor or medium with the solvent only. The concentrations used for each of the inhibitors are listed in table 1. Cells were inoculated with an even mixture of the three fluorescent recombinants for one hour at 4^o^C. The excess viruses were removed, and new medium containing the inhibitor (or solvent only) was added to cells. The infected cells were incubated at 37^o^C for 6 to 8 hours until images of the infected cells were taken. All inhibitors were tested for cell toxicity after 12 hours of incubation. In working concentration, none of the HDACi’s induced more than 5% cell death (tested by Propidium iodide positive cells). In 4 fold higher concentration, only Valporic Acid (VPA) had less than 80% viability. Anacardic Acid (AA) working concentration resulted in ~10% cell death and up to 40% in 4 fold higher concentration.

**Table 1.** List of inhibitors used in this work.

| **Inhibitor** | **Company** | **Working concentration** |
| --- | --- | --- |
| Trichostatin A (TSA) | Sigma Aldrich | 1.32μM |
| Suberohydroxamic Acid (SBX) | Sigma Aldrich | 40μM |
| Valporic Acid (VPA) | Merck | 4mM |
| Suberoylanilide Hydroxamic Acid (SAHA) | Sigma Aldrich | 10μM |
| Anacardic Acid (AA). | Merck | 250μM |

### Image acquisition and analysis

To estimate the number of HSV-1 genomes expressed in each infected cell, we obtained images (as described above) using a Nikon Eclipse Ti-E epifluorescence inverted microscope. Each experimental condition (different cells, viruses, inhibitors and MOI) was replicated in two wells, and the experiment was performed at least twice. From an individual well, five random areas were imaged. From each image, 100 cells were analyzed for their color content. To define the average number of incoming genomes being expressed, we used the mathematical equation for estimating the most likely average number of genomes expressed in each cell (λ) according to the number of one- (r1), two- (r2), or three-color (r3) cells out of the number of cells analyzed (n), as was previously developed (Kobiler et al., 2010):

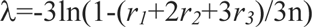

The λ was calculated for each well individually (based on 500 cells) and for each condition the mean λ and standard deviations were calculated. To predict the significance of the difference between the conditions a two tailed student T-test was performed.

### Live cell imaging

Vero cells plated in 8 well Nunc™ Lab-Tek™ II Chambered Coverglass were used. Infection with OK11 was carried out in the presence or absence of 1.32µM TSA as described above. Images were acquired using a Nikon Eclipse Ti-E epifluorescence inverted microscope every 10 min with DAPI and RFP fluorescence at 37^o^C in a 5% (vol/vol) CO2 enriched atmosphere using a Chamlide TC stage top incubator system (Live Cell Instrument). Two experiments were done with technical repeats (wells) per condition. From each well, five frames were taken and analyzed. First, using the Imaris 8.1 (Bitplane) image software we identified the individual cells according to the Hoechst DNA staining. The level of red fluorescence at each time point was measured for each identified cell by the software. We removed all cells in which fluorescence levels did not increase above 10% during the infection, as most of these cells were either dead or resistant to infection. From each well, at least 200 cells were analyzed. The average levels of fluorescence from all the cells analyzed per experiment were normalized, and the average and standard deviation between experiments are presented.

### Quantitative PCR

Freshly seeded cells were incubated in the presence or absence of 1.32µM TSA for three hours at 37^o^C. The entire RNA was extracted from the cells with the BIO TRI RNA (Bio-Lab ltd.) according to manufacturer protocol. The RNA was converted to cDNA with the High Capacity cDNA transcription kit (Applied Biosystems™) according to manufacturer protocol. cDNA was amplified using Faststart universal SYBR Green Master (Roche) in the presence of specific sets of primers for each gene (Table 2). qPCR was carried out and analyzed in StepOne™ (Applied Biosystems™). mRNA levels were normalized to the expression of GAPDH gene. The results were collected from five independent repeats. We performed an outlier identification using the extreme studentized deviate (EMD) method. One outlier result was removed for PML mRNA levels at HFF cells which did not change the significance of the results.

**Table 2.**
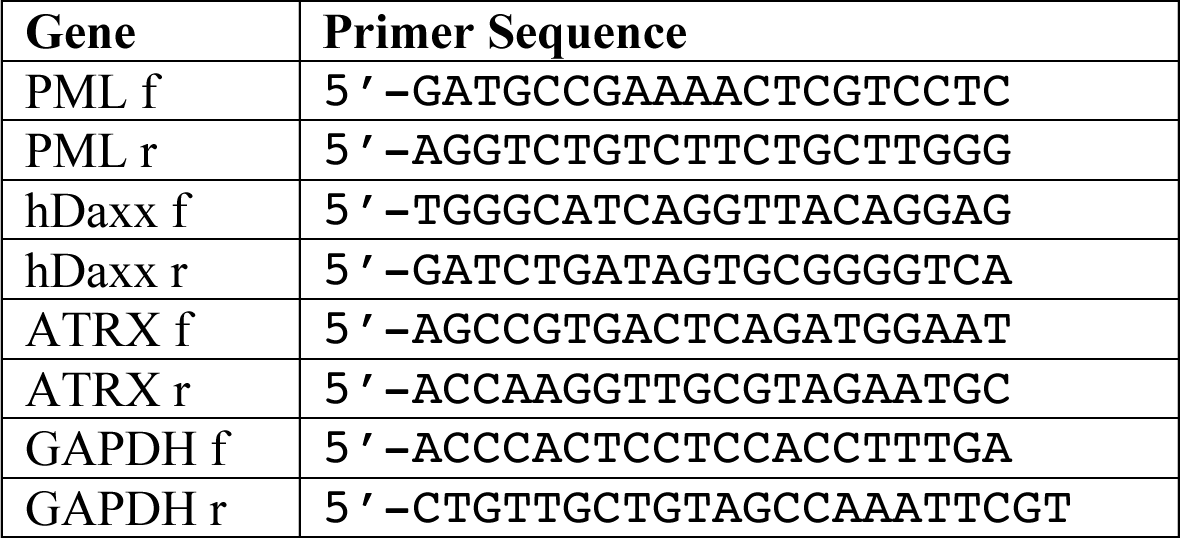
List of primers used for qPCR. Both forward (f) and reverse (r) primers for amplifying the specific genes are listed.

## Results

### HDACi treatment decreases the number of viral genomes establishing infection per cell

To study the role of histone modification on the number of viruses that initiate expression and replication in individual cells, we infected Vero cells in the presence or absence of several HDACi with three viral recombinants, each carrying a different fluorescent protein. We tested the following HDACi: Trichostatin A (TSA) - inhibits the class I and II HDACs, Suberohydroxamic Acid (SBX) – a competitive inhibitor that inhibits class II HDACs, Valporic Acid (VPA) – inhibitor of the class I HDACs and Suberoylanilide Hydoxamic Acid (SAHA) - inhibits the class I and II HDACs. All the HDACi were introduced three hours prior to infection. Figure 1A-D show representative images obtained 6-8HPI from such experiments. Surprisingly, incubating the cells with either one of the HDACi resulted in a lower average number of viral genomes expressed per cell (figure 1E). TSA, SBX and VPA all significantly decreased the average number of viral genomes expressed per cell at MOI 50 and 100, while with SAHA the decrease was not significant. (SAHA treatment decrease was significant in paired experiments compared to controls, data not shown). These results are unexpected, as HDACi generally increases open chromatin status and thus should promote higher expression from viral genomes. Thus, it is more likely that under the experimental conditions, the HDACi inhibitors are not affecting the chromatinization status of the viral genomes directly but probably affecting hosts' gene expression.

**Figure 1.**
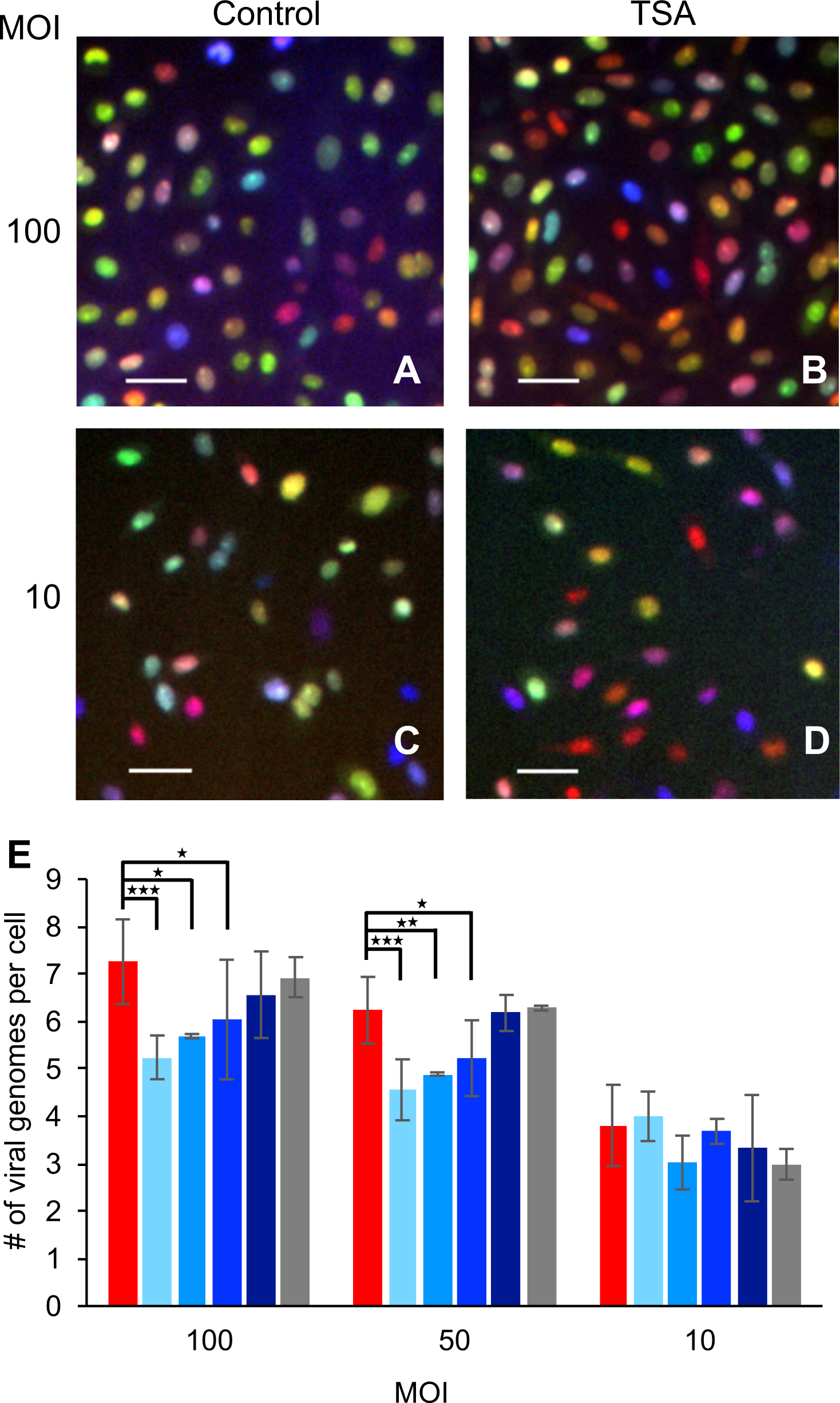
Effect of histone modifying enzyme inhibitors on the number of viral genomes expressed per cell. Vero cells were infected at MOI 100, 50 or 10 with a mixture of the three fluorophore expressing HSV-1 strains. (A-D) Representative images taken 6-8 hpi of infected cells at MOI 100 (A, B) or 10 (C, D) show the variability in the hues as a result of the three different fluorescent colors (mCherry –red, YFP – Green and mTurq2 – blue). Infection was carried out in the presence (B, D) or absence (A, C) of TSA. Scale bars 50µM. (E) Images were used for calculating the average number of genomes expressed per cell. The infected cells were incubated with different inhibitors, 3 hours prior to infection. Color coded as follows: No inhibitor - red, TSA - lightest blue, SBX - light blue, VPA – blue, SAHA – dark blue and AA in gray. Each bar represents at least two biological repeats and error bars represent standard deviations between the repeats. \**P* < 0.05, ** *P* < 0.01; \*\*\**P* < 0.001; by *t* test.

To ensure that our results are specific to HDACi, we also tested the effect of the histone acetyltransferases (HAT) inhibitor, Anacardic Acid (AA). In the presence of AA, no significant change in the number of expressed viral genomes per cell was observed in all MOI tested (figure1E). Our results show that all four HDACi tested, similarly, reduce the number of viral genomes expressed while the HAT inhibitor did not; suggesting that this phenomenon is dependent on the specific inhibition of the deacetylase activity.

### The viral progeny levels are independent of the number of genomes replicating per cell

To test the effect of the number of genomes replicating on viral progeny, we carried out single step growth curves in the presence or absence of the HDACi’s. We found that HDACi’s have very minor effect on the total outcome of infection according to the single step growth curves (figure 2A). On the other hand, the addition of AA resulted in a reduction of ~five folds compared to the control (P. value=0.018). Taken together these results indicate that there is no correlation between the number of viral genomes replicating, and the final concentration of progeny viruses released from cells.

**Figure 2.**
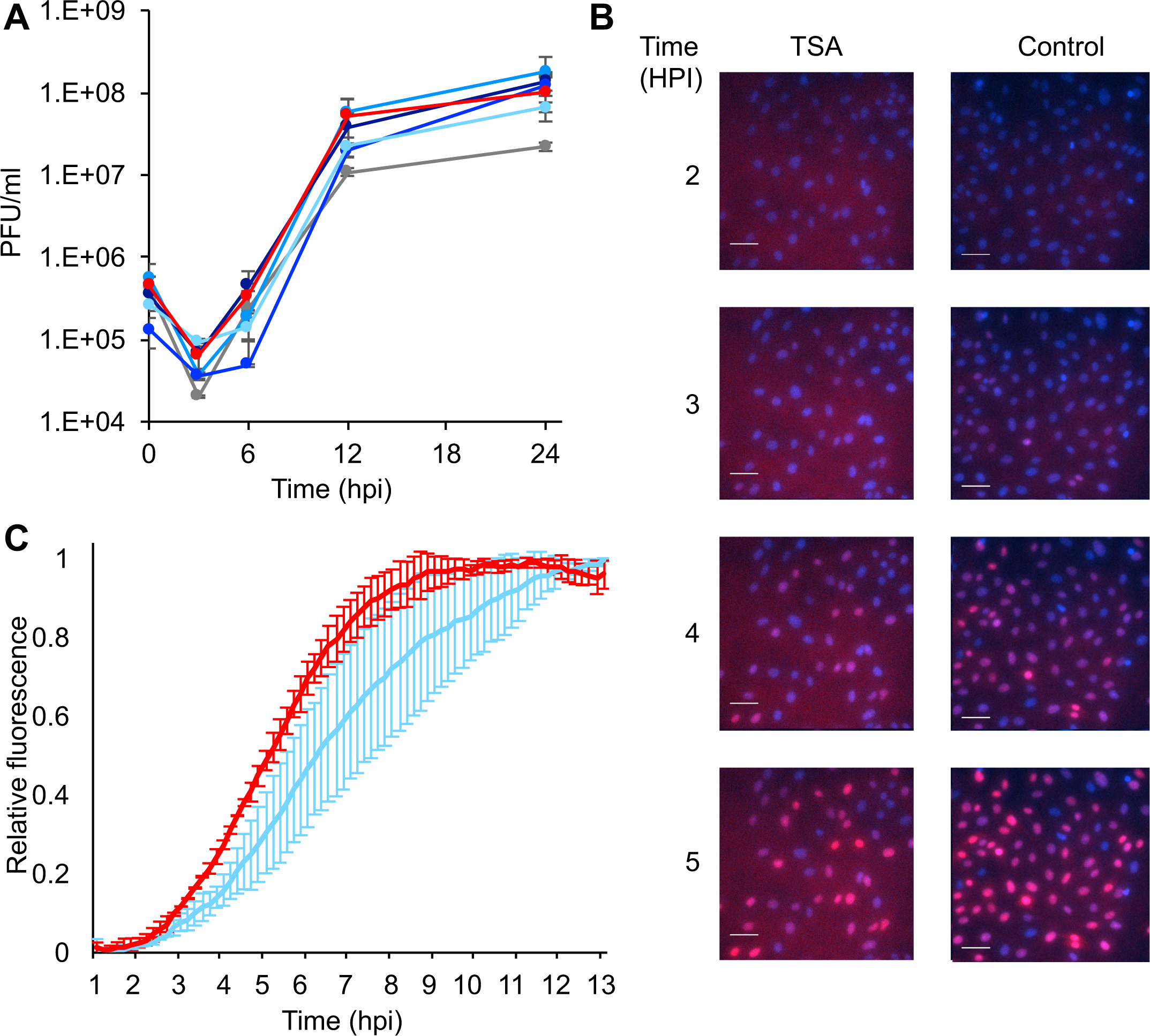
Effect of histone modifying enzyme inhibitors on the viral growth kinetics. (A) Vero cells were infected with an even mixture of the three fluorescent expressing HSV-1 recombinants at MOI 10 in the presence or absence of different inhibitors, and at different time points post-infection, progeny viruses were collected and assayed for titer. The infected cells were incubated with different inhibitors (color coded as in figure 1), 3 hours prior to infection. Each point is an average of viral titers obtained from three technical replicates. Error bars show standard deviations. (B-C) Vero cells were infected with an mCherry expressing recombinant at MOI 100 in the presence or absence of TSA. The cells were visualized every 10 minutes from one-hour post infection. (B) Representative images at the indicated time-points are shown. Cell nuclei were stained with Hoechst (Blue) and mCherry expression from the virus was monitored (red). Scale bars 50µM. (C) Each line represents the average relative fluorescence accumulation from two different experiments (color coded as in figure 1). In each experiment two wells were monitored (technical repeats), in each well 3-5 frames were analyzed. A total of more than 200 individual cells were collected from each well. Error bars show standard deviations between the two experiments.

Interestingly, both TSA and VPA showed a slower start compared to the untreated sample, i.e. almost no increase in viral progeny between 3 and 6 hours post infection. To test if HDACi have an effect on the kinetics of the viral infection, we monitored the infection using live cell microscopy. Vero cells were infected at MOI 100 with mCherry-expression viral recombinant in the presence or absence of TSA (Figure 2B). Figure 2C shows a slower accumulation of the mCherry signal in the presence of TSA, indicating slower kinetics of viral gene expression in TSA treated cells. These results corroborate the findings from the growth curve experiments and are in agreement with the finding that a lower number of genomes initiate expression in the presence of HDACi.

### Differences in the number of viral genomes expressed in human cells

As HSV-1 coevolved with its human host, it is likely that some host factors, that are involved in viral replication, might be specific to human cells (Lou et al., 2016). To test if the number of viral genomes replicating per cell is cell type dependent, we have infected two human cells types: the human foreskin fibroblasts (HFF) and U2OS cells (human osteosarcoma cell line). HFF cells are immortalized primary fibroblasts that are not known to have any cellular abnormalities. U2OS on the other hand, are known to be more permissive for HSV-1, VP16 and ICP0 mutant strains as U2OS cells are defective in several intrinsic-immunity cellular mechanisms (Yao and Schaffer, 1995; Smiley and Duncan, 1997; Hancock et al., 2006). For example, in U2OS cells, both copies of the ATRX gene are deleted (Heaphy et al., 2011). We tested the number of herpes genomes expressed per cell in HFF and U2OS cells. We found that in general, fewer viral genomes can initiate expression in HFF compared to Vero or U2OS, especially in the lowest MOI tested (10, Figure 3A and Table 3). In U2OS cells more viral genomes per cell are capable of establishing infection, with the biggest difference seen in the highest MOI 100 (figure 3A and Table 3). These results suggest that cellular factors are key regulators determining the number of viral genomes being replicated per cell.

**Figure 3.**
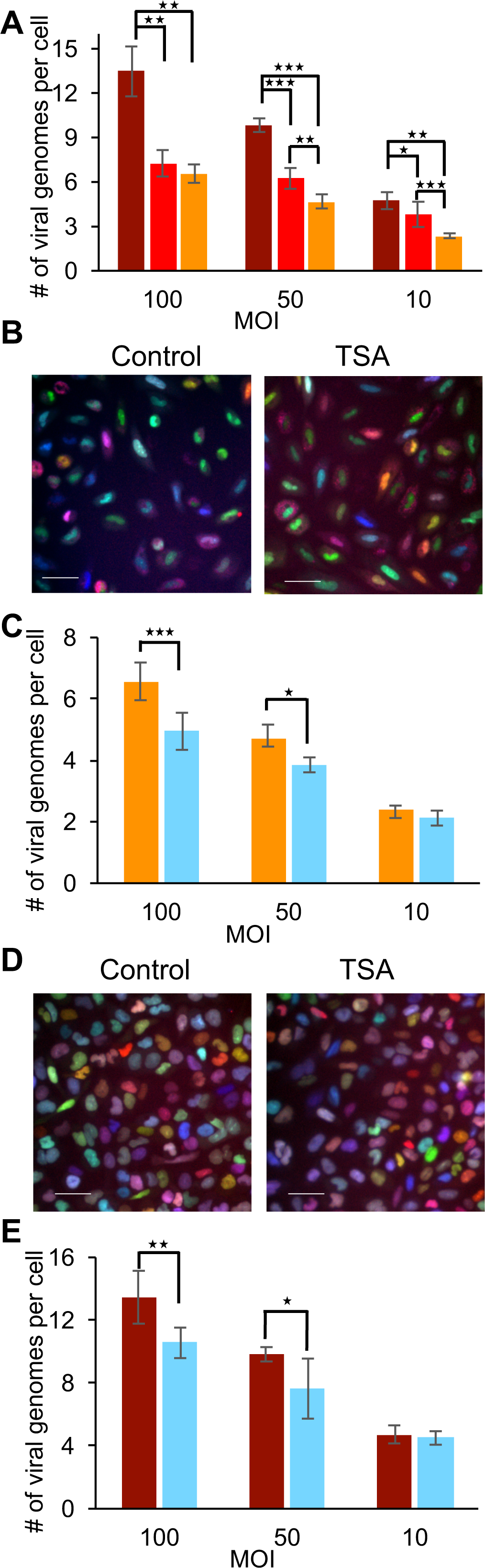
TSA reduces the number of viral genomes expressed per cell in human cell lines. **(A)** Two human cells lines: U2OS (Dark red) and HFF (Orange) were infected with a mixture of the three fluorophore expressing HSV-1 strains. The average number of viral genomes expressed in the human cells was compared to Vero (red) cells. HFF cells (B and C) or U2OS cells (D and E) were infected in the presence or absence of TSA with a mixture of the three fluorophore expressing HSV-1 strains. Representative images taken 6-8 HPI of HFF (B) and U2OS (D) cells infected at MOI 100 are shown. Scale bars 50µM. Images were used for calculating the average number of genomes expressed per cell. The average number of genomes in HFF cells (C) or U2OS cells (E) in the presence (light blue) or absence (HFF – orange and U2OS - dark red) of TSA at different MOI as indicated are presented. The infected cells were incubated with TSA, 3 hours prior to infection. Each bar represents six biological repeats and error bars represent standard deviations between the repeats. *P < 0.05, ** P < 0.01; ***P < 0.001; by t test.

**Table 3.** Summary of the average number of genomes expressed per cell under different conditions. The average and standard deviation of the number of genomes expressed per cell under different conditions as specified in the table was calculated.

|  | HFF |  | Vero |  | U2OS |  |  |
| --- | --- | --- | --- | --- | --- | --- | --- |
| TSA | + | - | + | - | + | - | ΔICP0 |
| MOI 100 | 4.9±0.6 | 6.6±0.6 | 5.2±0.5 | 7.3±0.9 | 10.6±1.0 | 13.5±1.7 | 7.4±0.4 |
| MOI 50 | 3.9±0.2 | 4.7±0.5 | 4.6±0.6 | 6.2±0.7 | 7.6±1.9 | 9.8±0.5 | 5.7±0.4 |
| MOI 10 | 2.1±0.2 | 2.4±0.2 | 4.0±0.5 | 3.8±0.9 | 4.5±0.4 | 4.7±0.6 | 2.8±0.4 |

### TSA treatment reduces the number of viral genomes initiating expression in human cells

Several studies have shown that the effect of HDACi on herpes replication is cell type dependent (Cody et al., 2014). To test if HDACi decreases the number of viral genomes in the human cell lines, we infected HFF and U2OS cells in the presence or absence of TSA (the HDACi with the highest effect on Vero cells). We found that the presence of TSA significantly reduces the number of viral genomes replicating per HFF cell in both MOI of 50 and 100 (Figure 3B and 3C). At MOI of 10 the reduction was not statistically significant. As with the other cell types in the presence of TSA, U2OS cells demonstrated a significant decrease in the number of viral genomes expressed per cell, both in MOI of 50 and 100 (Figure 3D and 3E). Interestingly, in all cells tested, infection in the presence of HDACi results in a similar degree of decline in the number of herpes viral genomes expressed per cell, regardless of the different basal levels of genomes establishing infection per cell. In the three cell types, infection in the presence of TSA results in a reduction of ~25% at an MOI 100 and 10% or less in MOI10 (Table 3). Taken together, these results suggest that the effect HDACi have on the viral initiation of replication is independent of cell type.

### Treatment with HDACi increases the levels of antiviral ND10 proteins

The ability of HDACi to decrease the number of viral genomes initiating expression per cell is contra-intuitive to the predicted role of HDACi on viral genomes. As we observed significant differences among the cell lines tested (figure 3A and table 3), we assume that alterations in the levels of host factors influence the probability of viral genomes to initiate expression and replication. We hypothesize that the treatment with HDACi results in higher expression of cellular genes that are involved in inhibiting viral replication. To test this hypothesis, we selected three genes, PML, ATRX and hDaxx, all part of the ND10 bodies and known to be involved in the intrinsic response against HSV-1. We tested the mRNA levels of these genes following three-hour incubation in the presence or absence of TSA. In HFF cells, the mRNA levels of ATRX were significantly upregulated in the presence of TSA (P<0.05), whereas both PML and hDaxx mRNA levels were upregulated to a lesser degree (figure 4). In U2OS cells, there is no ATRX mRNA and the PML mRNA levels increased significantly (P<0.05) due to the TSA treatment. While the increase of PML mRNA levels in U2OS was higher than in HFF, hDaxx mRNA levels did not change in the presence of TSA in U2OS. Our results suggest that some host antiviral genes are overexpressed in the presence of TSA. We speculate that the higher levels of antiviral genes expression following treatment with HDACi results in a hostile environment for the entering viral genomes, reducing the number of genomes initiating expression and replication.

**Figure 4.**
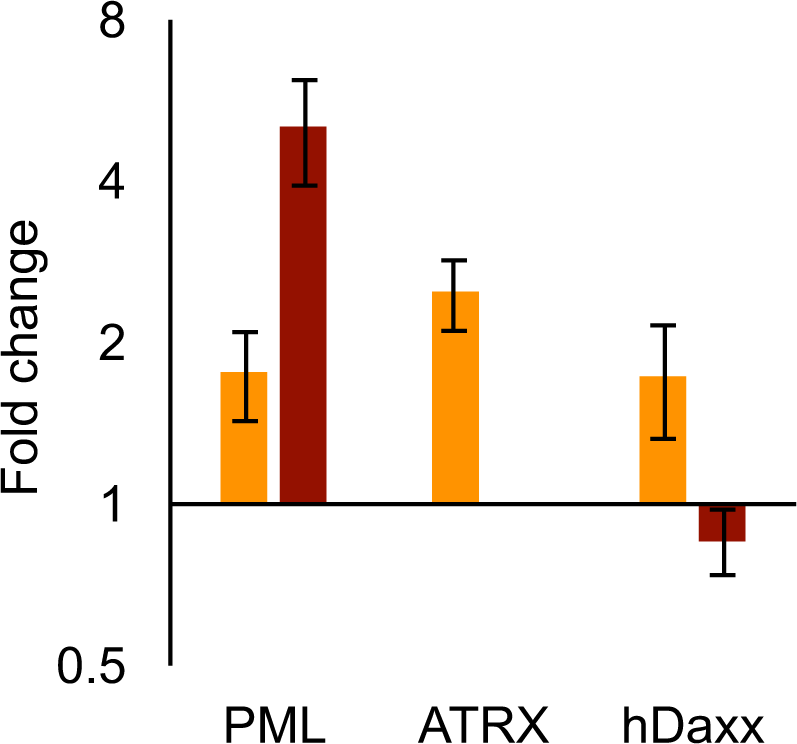
TSA treatment of HFF and U2OS cells increases intrinsic immunity gene expression. qRT-PCR measurements of host genes (as marked) following incubation of HFF (Orange) or U2OS (dark red) cells in the presence or absence of TSA. Each bar represents the mean difference between TSA treatment and control in five independent experiments, each experiment was tested in duplicates. Error bars represent standard errors of mean.

### ICP0 deletion reduces the number of viral genomes initiating expression in U2OS cells

Many host genes, including PML and ATRX, that have a role in the anti-herpes intrinsic response are counteracted by the viral ICP0 protein. To test the role of ICP0 in determining the number of viral genomes that establish infection per cell, we first constructed three viral recombinants (each carrying a different fluorescent protein) into the dl1403 (ICP0 null) background (see material and methods). All three new ICP0 deleted fluorophore expressing HSV-1 strains: OK29 (mCherry) OK32 (mTurq2) and OK40 (EYFP), show similar single cell growth curves (figure 5A). While the efficiency of plating of ICP0 positive viral strain on Vero and U2OS cells is similar (ratio is close to one), the ICP0 deleted strains are forming more plaques on U2OS cells (known to be permissive for ICP0 null mutants) as expected (figure 5B). U2OS cells were infected with an even mixture of all three ICP0 null recombinants at different MOIs. In the absence of ICP0, a significant decrease in the number of viral genomes that initiate expression compared to wild type infection can be detected in all MOIs tested (figure 5C and Table 3). This decrease is almost two folds and results in numbers similar to those obtained by wild type infection in HFF cells. (We were unable to obtain titers for the ICP0 null recombinants that will be sufficient for HFF infection at comparable MOIs). Our findings indicate that the ICP0 protein has a major role in allowing viral genomes to initiate expression, even in U2OS cells, probably by promoting the degradation of antiviral host factors.

**Figure 5.**
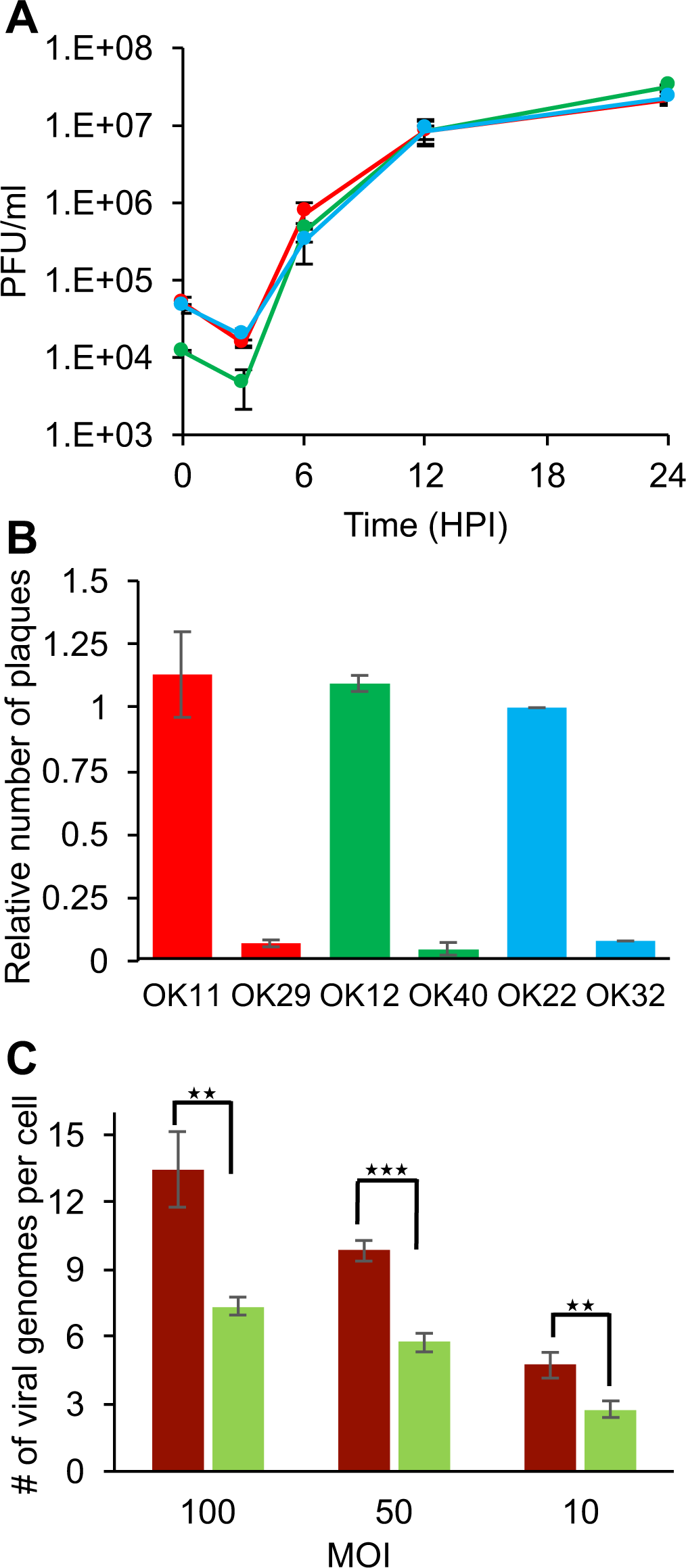
Deletion of the ICP0 gene reduces the number of viral genomes expressed per cell in U2OS cells. (A) U2OS cells were infected with one of the ICP0 deleted fluorophore expressing HSV-1 strains: OK29 (red) OK32 (blue) and OK40 (green). At different time points post-infection, progeny viruses were collected and assayed for titer. Each point is an average of viral titers obtained from three technical replicates. Error bars show standard deviations. (B) The titers of fluorophore expressing HSV-1 strains (either ICP0 plus strains: OK11, OK12 and OK22 or ICP0 deleted strains: OK29, OK40 and OK32) were measured on Vero cells or U2OS cells. The ratio between titers on Vero cells to U2OS cells is presented for each strain as marked. Each bar represents the ratio between two biological repeats and error bars represent standard deviations between the repeats. (C) U2OS cells were infected with a mixture of the three ICP0 deleted fluorophore expressing HSV-1 strains (light green). The average number of viral genomes expressed was compared to wild type infection of U2OS cells (Dark red). Images taken 6-8 hpi were used for calculating the average number of genomes expressed per cell. Each bar represents six biological repeats and error bars represent standard deviations between the repeats. ** P < 0.01; ***P < 0.001; by t test.

## Discussion

We have previously shown that only a finite number of herpesvirus genomes can establish infection in a given cell (Kobiler et al., 2010; Kobiler et al., 2011; Taylor et al., 2012). Several other viruses from different families, including HIV and plant RNA viruses, were also shown to have a restricted number of genomes initiating expression per cell (Gonzalez-Jara et al., 2009; Gutierrez et al., 2010; Miyashita and Kishino, 2010; Del Portillo et al., 2011), suggesting a general characteristic in viruses' life cycles. Here, we identified that the HDACi reduces the number of genomes expressed in a single cell in three different cell types. We speculated that HDACi activity on host chromatin disrupts the balance between the virus and host cell, by allowing higher expression of intrinsic immunity proteins. In support of this hypothesis, we found that U2OS cells which are known to be defective in their intrinsic immunity, express greater numbers of viral genomes compared to the other cell types. Moreover, HDACi treatment, which reduces the number of genomes initiating gene replication, induces the expression of cellular proteins involved in inhibition of viral functions.

We consistently observed that treatment with HDACi reduce the number of parental viral genomes initiating expression. These results were observed in the presence of different HDACi’s (figure 1E) and for three different cell types (figures 1E and 3C, E). However, a significant decrease in the number of viral genomes was only observed in MOI 100 and 50. We found that at MOI 10, while a small decrease is observed (in most cases), a statistically significance is not obtained at this MOI. We previously suggested that at MOI 10 and lower the effect of the incoming viruses is more significant in determining the number of viral genomes, expressing a higher percentage of incoming viruses able to establish infection compared with the higher MOIs (Kobiler et al., 2010). Thus, in higher MOIs, cell factors probably have a larger effect on the number of genomes initiating expression. Our results suggest that the HDACi effect on incoming genomes is mediated by overexpression of host antiviral proteins (figure 4 and 5C). Taken together, this might explain why no significant effect of HDACi on viral genome numbers is observed at MOI 10.

Our results indicate that there is no correlation between the average number of genomes that initiate expression to the progeny outcome from the infection (compare figures 1 and 2A). This finding raises the question: what is the biological advantage of regulating the number of viral genomes initiating expression and replication per cell? One possible mechanism involves slowing down the rate of the infection process. Our results support this idea as treatment with HDACi reduces the number of viral genomes and decreases the rate in which the viral infection progresses (Figure 2B). Similarly, when lowering the MOI (in MOI higher than three where most cells are infected), there is no significant effect on the progeny outcome however there is a reduction in the rate in which the infection proceeds. Corroborating this observation, infection with higher MOI results in higher average number of genomes expressed per cell. Thus, the evolutionary advantage of reducing the quantity of viral genomes that initiate replication may be to prevent cell death before viral particles can be released.

Another possible evolutionary advantage is the maintenance of viral fitness and diversity. Recently it was shown that tomato mosaic virus, a positive-strand RNA virus, by random selection of a limited number of genomes initiating replication in infected cells, is able to selectively propagate more advantageous genotypes (Miyashita et al., 2015). We speculate that this mechanism can also be also important for DNA viruses, as much of their diversity is governed by recombination (Szpara et al., 2014). The methods presented here to modify the number of viral genomes could provide ways to identify the evolutionary advantage of replicating only a finite number of genomes in individual cells.

We identified significant differences between the three cell types we tested (Figure 3A). We found that in U2OS cells, more incoming viral genomes are expressed compared to HFF cells. In Vero cells, the number of genomes was higher than those of HFF but significantly lower than those of U2OS cells. It is important to note that Vero cells originated from green monkey whereas both HFF and U2OS are of human origin. HFF, human diploid cells have both the intrinsic and the innate immune system functioning. Vero cells are incapable of producing type I interferon as many of the type I interferon genes are missing because of a ~9Mb deletion on chromosome 12 (Osada et al., 2014); thus they have a dysfunctional innate immunity. U2OS do not express ATRX and have lower expression of other intrinsic immunity proteins (Lukashchuk and Everett, 2010), thus exhibiting impaired intrinsic immunity and facilitating genomes expression. We suggest that the intrinsic immunity has a bigger role than the innate immunity in repressing incoming viruses expression. In agreement with this hypothesis, we found that the HDACi treatment induces the expression of PML and ATRX, which are part of the intrinsic immunity system (figure 4).

The genes tested here, PML, ATRX and HDAXX are part of the nuclear domains -ND10 (also known as PML bodies) that among other functions are involved in recognizing and inhibiting viral genomes in the nucleus (Glass and Everett, 2013; Xu et al., 2016). ND10 bodies are specifically known to interact with HSV-1 genomes and be dismantled by the viral ICP0 protein (Everett and Maul, 1994; Maul and Everett, 1994; Boutell and Everett, 2013). ICP0 deletion recombinants show a significant decrease in the number of genomes expressed per cell (Figure 5C). Taken together these results indicate that perturbation of the viral host interactions balance by either inducing the expression or increasing the stability of host genes can both reduce the probability of a viral genome to establish infection.

The response to HDACi treatment is expected to result in higher gene expression, however prior studies have identified that while many genes are over-expressed following HDACi treatment, innate immunity genes are down regulated (Nusinzon and Horvath, 2003; Suh et al., 2010). Thus, the increase in expression of PML and ATRX we observed following HDACi treatment may reflect a specific regulation of this part of the immune response. Interestingly, inhibition of HDAC by TSA dramatically enhanced induction of antimicrobial peptides but not of proinflammatory cytokines (Fischer et al., 2016), further indicating selective expression post HDACi treatment of different parts of the immune system.

In conclusion, this work provides evidence for viral host interactions in regulating the number of viral genomes establishing infection per cell. We suggest that inhibition of HDAC during infection can be more significant for the host chromatin than to the viral chromatin. The number of incoming viral genomes that initiate expression and replication is probably dependent on the intrinsic immunity state of the infected cell and influence the infection process but has a little effect on the number of progeny viruses.

## Acknowledgments

We thank Sara Selig for the HFF cells and Roger Everett for the dl1403 HSV-1 mutant. We thank all members of the Kobiler lab for valuable discussions in the preparation of this manuscript.

